# *Listeria monocytogenes* Co-opts Caveolin-Mediated E-cadherin Trafficking and Macropinocytosis for Epithelial Cell-to-Cell Spread

**DOI:** 10.1101/2022.04.06.487361

**Authors:** Prathima Radhakrishnan, Mugdha Sathe, Julie A. Theriot

## Abstract

*Listeria monocytogenes* is an intracellular bacterial pathogen that spreads directly between adjacent host cells without exposure to the extracellular space. Recent studies have identified several host cell factors that promote *L. monocytogenes* cell-to-cell spread in epithelial monolayers, but details of the mechanism remain unclear. We find that the adherens junction protein, E-cadherin, promotes *L. monocytogenes* cell-to-cell spread at the recipient side of cell contacts. In particular, two point mutations in E-cadherin’s cytoplasmic domain that prevent its ubiquitination hinder bacterial cell-to-cell spread efficiency without reducing the extent of contact between neighboring cells. As ubiquitination induces E-cadherin endocytosis, we hypothesize that E-cadherin promotes protrusion engulfment, where donor cell protrusions containing *L. monocytogenes* are taken up by the recipient cell concurrently with E-cadherin internalization. In support of this hypothesis, we show that inhibiting caveolin-mediated membrane trafficking reduces *L. monocytogenes* cell-to-cell spread only under conditions where E-cadherin can be ubiquitinated. Additionally, we demonstrate that macropinocytosis also contributes to dissemination of *L. monocytogenes* through an epithelial monolayer.

## INTRODUCTION

*Listeria monocytogenes* is a facultative bacterial pathogen that causes spontaneous abortions and illness in immunocompromised individuals, ranking third among foodborne pathogens in number of deaths caused per year (Hamon et al., 2006). When ingested orally, *L. monocytogenes* first infects epithelial cells of the human gut (Schlech and Acheson, 2000). It adheres to these cells by binding receptors such as E-cadherin on the host cell surface (Mengaud et al., 1996, Lecuit et al., 1999, Ortega et al., 2017) and is then actively taken up by the host cells. Upon entry, the bacterium secretes phospholipases and the pore-forming toxin Listeriolysin O (LLO) to rupture its vacuole and escape into the host cell’s cytoplasm (Portnoy et al., 1988, Marquis et al., 1995). It propels itself within the cytoplasm using actin-based motility (Tilney and Portnoy, 1989, Dabiri et al., 1990, Theriot et al., 1992), eventually reaching the plasma membrane and forming a membrane-covered protrusion that can push into the neighboring cell. Intercellular spread occurs when the recipient cell engulfs the *L. monocytogenes*-containing protrusion and the bacterium escapes the secondary vacuole into the cytoplasm of the recipient cell (Robbins et al., 1999). This process allows the pathogen to propagate through monolayers without exposure to the extracellular space, thereby avoiding the host’s humoral immune response (Zenewicz and Shen, 2007).

Several molecular factors have been identified in both the pathogen and the host cell that promote the formation or elongation of protrusions at the donor side of cell contacts and enhance the efficiency of cell-to-cell spread. For example, the secreted bacterial virulence factor internalin C (InlC) supports protrusion formation and elongation by relieving cortical tension at the donor side of cell contacts (Rajabian et al., 2009) and recruiting exocyst machinery to deliver additional membrane to protrusions (Dowd et al., 2020). By secreting cyclic-di-AMP, *L. monocytogenes* enhances nitric oxide production in host cells, which increases the speed of the pathogen’s actin-based motility, to allow for the generation of protrusions that are more frequently taken up by a recipient cell (McFarland et al., 2018).

Two recent studies have identified specific host cell factors that participate in cell-to-cell spread at the recipient side rather than the donor side of cell contacts (Sanderlin et al., 2019, Dhanda et al., 2020). Using an RNAi screen to identify host cell factors that contribute to *L. monocytogenes* cell-to-cell spread, Sanderlin et al. identified caveolin-1, a mediator of caveolar endocytosis, as contributor to bacterial spread in recipient cells (Sanderlin et al., 2019). Using a more targeted candidate-based approach, Dhanda et al. also described a role for caveolar elements including caveolin-1, cavin-1, and EHD2 in promoting *L. monocytogenes* cell-to-cell spread, along with other endocytosis-related proteins including dynamin (Dhanda et al., 2020).

In addition to caveolins, the RNAi screen conducted by Sanderlin et al. also found a role for the epithelial cell adherens junction protein, E-cadherin, in promoting cell-to-cell spread (Sanderlin et al., 2019). E-cadherin is robustly detected all along protrusions containing *L. monocytogenes* (Robbins et al., 1999), and cadherin expression is required for the epithelial cell-to-cell spread of *Shigella flexneri*, another intracellular pathogen that uses actin-based motility for spread in epithelial monolayers (Sansonetti et al., 1994). Interestingly, adhesion proteins including VE-cadherin and nectins have been shown to promote trans-endocytosis, the process by which cytoplasmic material and transmembrane proteins are exchanged between neighboring cells through a receptor-mediated process (Cagan et al., 1992, Sakurai et al., 2014, Generous et al., 2019). For these reasons, we began this study of *L. monocytogenes* cell-to-cell spread by investigating the mechanism by which the adhesion protein, E-cadherin, promotes bacterial intercellular spread. Our results are consistent with a simple model where caveolin-mediated trafficking of E-cadherin at the recipient side of cell-cell contacts promotes internalization of *L. monocytogenes*-containing protrusions, and thereby contributes to spread.

## RESULTS

### The Cytoplasmic Domain of E-cadherin Promotes *Listeria monocytogenes* Cell-to-Cell Spread at the Recipient Side of Cell Contacts

To quantify the extent to which E-cadherin influences *L. monocytogenes* cell-to-cell spread, we compared efficiency of cell-to-cell spread in epithelial cell monolayers grown in tissue culture. To this end, we used A431D epithelial cells, a cell line derived from a human primary epidermoid carcinoma that have lost expression of E-cadherin but retain expression of accessory proteins associated with adherens junctions (Lewis et al., 1997). Because of this property, it is convenient to use A431D cells as the background for expression of E-cadherin constructs with specific molecular lesions, in order to evaluate the roles of the protein’s various functional domains in complex biological processes (McEwen et al., 2014). For our first measurement, we compared the efficiency of *L. monocytogenes* spread between A431D epithelial cells expressing full length (wild-type) E-cadherin (WT E-cad) and parental A431D cells expressing no E-cadherin (Null E-cad). As E-cadherin is required for the initial invasion step through which the bacterium enters the monolayer from the extracellular space (Mengaud et al., 1996, Lecuit et al., 1999), we compared focus size at 12 hours post-infection in a monolayer of WT E-cad cells with 1:100 WT E-cad: Null E-cad A431D cells (Fig. 1A-B, Suppl. Movie 1), expecting that focus size at this late time point would be dominated by spread between the numerous Null E-cad cells rather than the single WT E-cad cell required for initial invasion. Because *L. monocytogenes* foci in epithelial monolayers are highly irregular in shape (Ortega et al., 2019), simple methods for determining the size of the focus using the smallest convex hull typically overestimate the true extent of cell-to-cell spread (Suppl. Fig. S1A). As an alternative, we optimized a method for calculating a tightly wrapped “alpha-shape” (Edelsbrunner et al., 1983) that more accurately correlates with the total number of infected host cells in a focus (Suppl. Fig. S1B). Using this metric (Fig. 1C-D), we found that the focus area was diminished by about 50% when *L. monocytogenes* spread between Null E-cad cells as compared to WT E-cad cells (Fig. 1E), suggesting that E-cadherin facilitates but is not strictly required for *L. monocytogenes* cell-to-cell spread.

**Figure 1:**
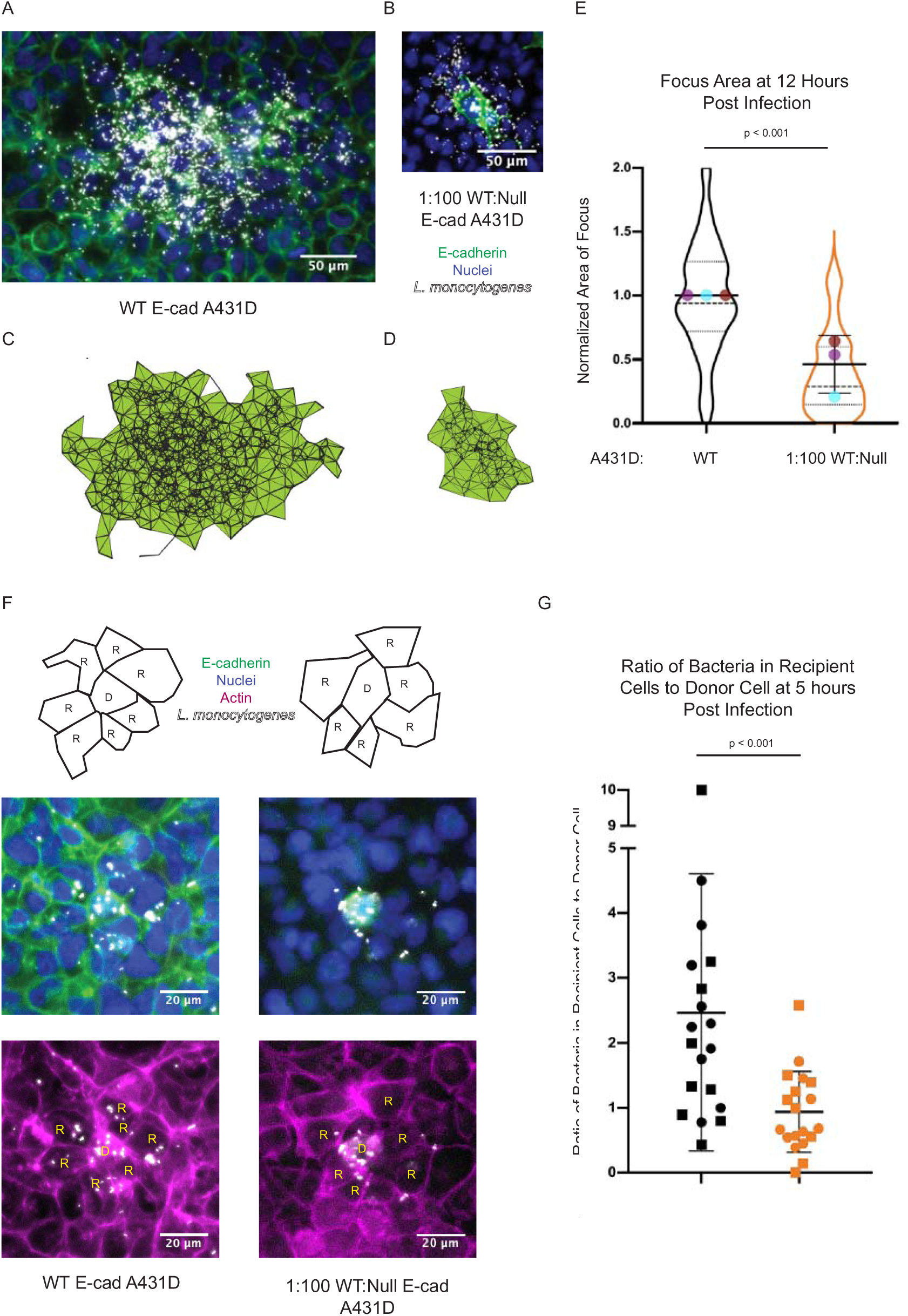
E-cadherin promotes *L. monocytogenes* cell-to-cell spread at the recipient side of cell contacts. **A.** *L. monocytogenes* focus in A431D epithelial cell monolayer expressing full-length GFP-tagged E-cadherin (WT E-cad), imaged at 12 hours post infection (h.p.i.). Bacterial fluorescence (mTagRFP) is shown in white, GFP-E-cad in green, and DAPI (labeling host cell nuclei) in blue. **B.** *L. monocytogenes* focus in a mixed monolayer with a ratio of 1:100 WT E-cad to Null E-cad cells, imaged at 12 h.p.i. Colors as in part A. The single GFP-E-cad cell that was the initial site of infection is visible at the center of the field of view. **C.** Alpha shape corresponding to the bacterial focus shown in part A. **D.** Alpha shape corresponding to the bacterial focus shown in part B. **E.** Distribution of focus sizes at 12 h.p.i. for *L. monocytogenes* in WT E-cad host cells (left) and in mixed monolayers (right). Colored dots represent the mean alpha shape area for three independent experiments, normalized to the control (WT E-cad condition) on each day. Violin plots show the full range of focus sizes, with 59 total foci for each condition. Thick black line shows the mean, whiskers show the standard deviation, dashed line shows the median, and dotted lines the first and third quartiles. The p-value was determined using the linear mixed-effects model (see Materials and Methods). **F.** Schematic (top) and immunofluorescence images (bottom) representing the design of the experiment to measure the efficiency of bacterial transfer from a WT E-cad donor cell (D) to a group of WT E-cad or Null E-cad recipient cells (R). Center row shows *L. monocytogenes* fluorescence (white), GFP-E-cad (green) and DAPI (blue). Lower row shows F-actin (fluorescent phalloidin) for the same fields of view. **G.** Ratio of the sum of bacteria in all recipient cells to the number of bacteria within the initial donor cell for transfer from a WT E-cad donor to WT E-cad recipient cells (left) and for transfer from a WT E-cad donor to Null E-cad recipient cells (right). Each point represents an individual focus. Squares and circles represent independent experiments performed on two separate days. Horizontal lines show the means and whiskers show the standard deviation. The p value was determined using the Wilcoxon rank-sum test.

E-cadherin might promote spread by aiding in the formation of protrusions at the donor cell, uptake of protrusions at the recipient cell, or both. To distinguish between these possibilities, we infected WT E-cad cells or 1:100 WT E-cad: Null E-cad A431D cells for a short time period of five hours, such that only one round of cell-to-cell spread would have occurred. Then, for each focus, we quantified the ratio between the total number of bacteria in neighboring recipient cells to the number of bacteria in the originally infected donor cell, thereby comparing the efficiency of transfer from WT E-cad to WT E-cad cells with the efficiency of transfer from WT E-cad to Null E-cad cells (Fig. 1F). This ratio was significantly higher for spread into WT E-cad recipient cells (Fig. 1G), although the total number of bacteria in each focus was comparable for both conditions (Suppl. Fig. S1D). As the lack of E-cadherin in the recipient cell specifically reduced the efficiency of cell-to-cell spread, we conclude that E-cadherin is involved in the uptake of protrusions at the recipient side of cell contacts, although we cannot rule out the possibility that it might also participate in the formation of protrusions by the donor cell.

To further explore the mechanism by which E-cadherin promotes spread, we compared *L. monocytogenes* cell-to-cell spread through a monolayer of WT E-cad cells with spread through a monolayer of A431D cells expressing an E-cadherin truncation mutant that lacked its entire cytoplasmic domain, deleting amino acids 731-822 (Δcyto E-cad) (Ringwald et al., 1987). Although this truncated protein was enriched at cell-cell junctions, it was unable to recruit cytoplasmic binding partners such as β-catenin (Fig. 2A). Focus area was reduced in the Δcyto E-cad cells as compared to the WT E-cad cells, nearly to the extent that focus area was reduced in the Null E-cad cells (Fig. 2B). Using a strain of *L. monocytogenes* deficient for cell-to-cell spread, we confirmed that the initial replication rate over the first 3 to 5 hours post-infection for bacteria in WT E-cad cells and in Δcyto E-cad cells was indistinguishable (Suppl. Fig. S2A), consistent with the idea that these focus size defects are caused by a specific deficiency in cell-to-cell spread rather than effects on earlier events in bacterial invasion or replication. Our results suggest that it is not E-cadherin’s extracellular domain, which physically links neighboring cells (Shapiro et al., 1995), that is primarily involved in augmenting *L. monocytogenes* cell-to-cell spread. Rather, E-cadherin’s cytoplasmic domain, which reinforces cell-cell junctions due to its linkage to the underlying actin cytoskeleton (Buckley et al., 2014) and enables E-cadherin internalization and turnover (Delva and Kowalczyk, 2009), contributes to *L. monocytogenes* cell-to-cell spread.

**Figure 2:**
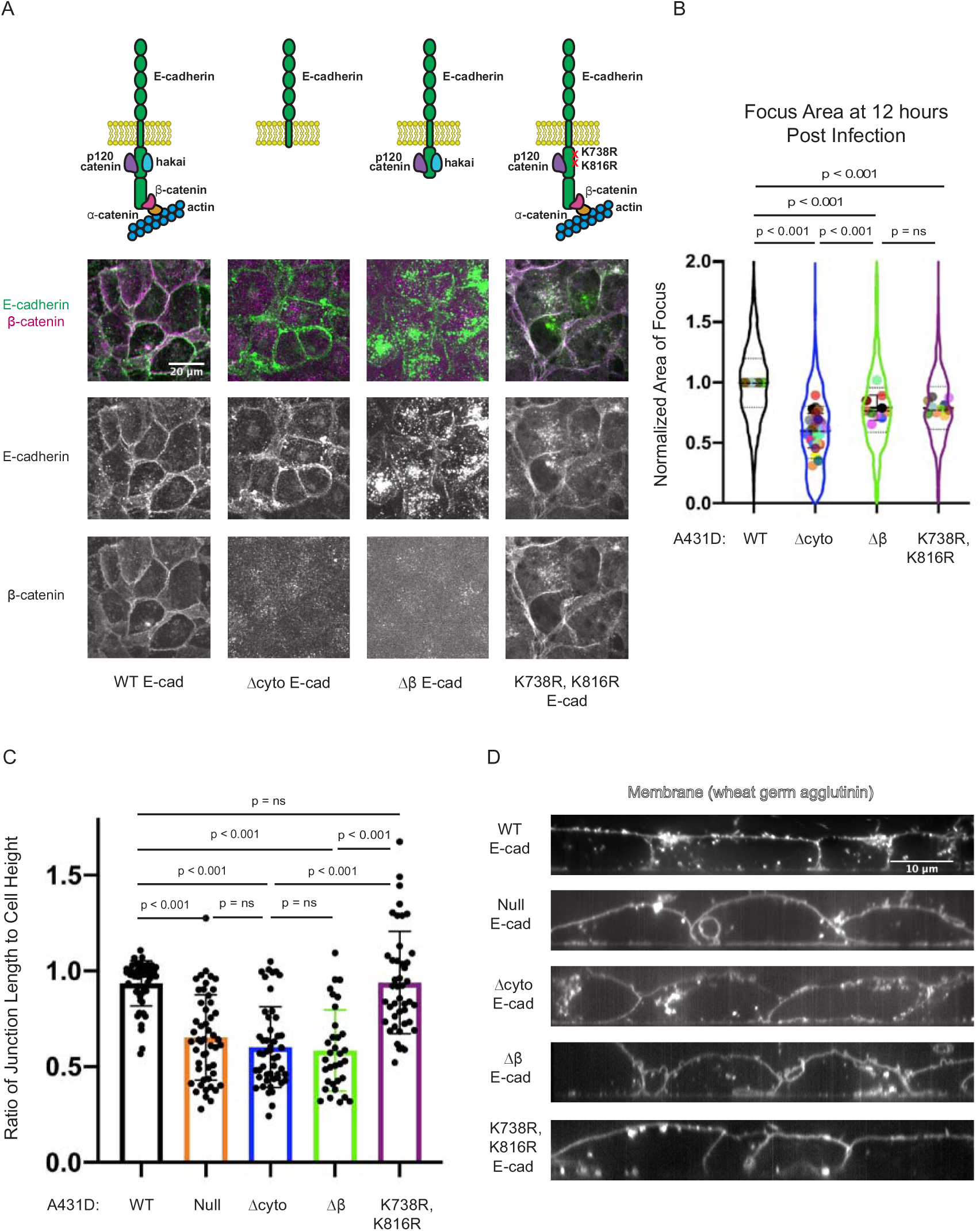
Mutations in E-cadherin’s cytoplasmic domain reduce efficiency of *L. monocytogenes* cell-to-cell spread. **A.** Schematic representations of WT, Δcyto, Δβ, and K738R, K816R E-cadherin (top) and immunofluorescence images of E-cadherin and β-catenin in these cells (bottom). Δcyto and Δβ E-cadherin localize to cell junctions but do not bind β-catenin. K738R, K816R E-cadherin localizes to cell junctions and binds β-catenin. **B.** Focus size for *L. monocytogenes* infection in Δcyto, Δβ and K738R, K816R E-cadherin-expressing host cells, normalized to focus size for infection of WT E-cadherin-expressing cells performed in parallel. Each colored dot represents a separate experimental day. Violin plots show the full range of focus sizes, thick black line shows the mean, whiskers show the standard deviation, dashed line shows the median, and dotted lines the first and third quartiles. Number of independent experiments imaged for each condition from left to right was 37, 26, 10, and 7. Number of individual foci imaged for each condition from left to right was 1979, 968, 551, and 435. P values were determined using the linear mixed-effects model. **C.** Ratio of junction length to cell height in WT, Null, Δcyto, Δβ and K738R, K816R E-cad cells. Between 33 and 50 junctions were analyzed for each cell line. Mean ± SD is shown on the graph and p values were determined using the Wilcoxon rank-sum test. **D.** Representative orthogonal views of membrane-labeled A431D cells from each of the five host cell lines.

In order to further narrow down the part of E-cadherin’s cytoplasmic domain that contributes to bacterial spread, we used a line of A431D cells expressing an E-cadherin truncation mutant (missing amino acids 810-882) that cannot bind β-catenin (Δβ E-cad) and therefore does not associate with the actin cytoskeleton, although it retains other protein-protein interaction sites in the juxtamembrane domain (JMD) (Ferber et al., 2002, Fujita et al., 2002) (Fig. 2A). Efficiency of bacterial cell-to-cell spread in this host cell line was intermediate between that in the WT E-cad cell line and the Δcyto E-cad cell line (Fig. 2B), suggesting that the lack of association between Δcyto E-cad and the actin cytoskeleton may contribute to the defect in bacterial cell-to-cell spread, but cannot completely explain the magnitude of this effect.

As β-catenin links E-cadherin to cortical actin (Yamada et al., 2005), cell junctions between neighboring Δβ E-cad or Δcyto E-cad cells are likely to be under less tension than cell junctions between neighboring WT E-cad cells (Verma et al., 2012). Moreover, Δβ E-cad or Δcyto E-cad cells would be unable to extend points of adhesion between two cells into larger zones of contact as this junctional reinforcement is normally a consequence of actin nucleation at cell junctions (Verma et al., 2004). The extent of cell-cell contact in an epithelial monolayer is expected to depend on the physical packing density of cells within the monolayer as well as on the ability of the cells to reinforce junctional integrity by attachment to the actin cytoskeleton. We therefore carefully controlled cell plating density to ensure that the physical packing of cells in the monolayer was consistent across all experiments (Suppl. Fig. S2B). In order to examine the extent of cell-cell contact more directly, we used structured illumination microscopy (SIM) to measure the height of junctions in the monolayers as compared to the maximum height of the cells (Suppl. Fig. S2C). Using this assay, we found that the extent of cell-cell contact between neighboring cells in a monolayer of A431D cells expressing WT E-cad was substantially greater than the extent of contact for Null E-cad cells plated at the same density, as expected (Fig. 2 C-D). Comparing the extent of cell-cell contact for both Δcyto E-cad and Δβ E-cad cell monolayers using this same assay, we found that neither truncated protein construct was able to rescue the size of cell-cell contacts beyond the extent observed for Null E-cad cells (Fig. 2 C-D). Therefore, the reduction of focus size of Null E-cad cells or Δcyto E-cad cells in comparison to Δβ E-cad cells cannot be explained by a physical inability of the bacterium to pass from donor to recipient cell simply because of a decreased area of cell-cell contact. Instead, we hypothesized that E-cadherin’s juxtamembrane domain (JMD), the portion that lies between the transmembrane domain and β-catenin binding domain, might specifically contribute to *L. monocytogenes* cell-to-cell spread.

### Preventing Ubiquitination of E-cadherin’s Juxtamembrane Domain Hinders *L. monocytogenes* Cell-to-Cell Spread

The E-cadherin JMD includes binding sites for several interaction partners that might contribute to *L. monocytogenes* cell-to-cell spread, including p120 catenin (Ferber et al., 2002) and the ubiquitin ligase hakai (Fujita et al., 2002). We constructed a modified full-length E-cadherin including two point mutations, K738R and K816R, that block ubiquitination and normal turnover of the E-cadherin JMD (Hartsock and Nelson, 2012), and expressed this construct in A431D cells. In contrast to Null E-cad cells, Δcyto E-cad cells, and Δβ E-cad cells, monolayers consisting of K738R, K816R E-cad cells exhibited normal recruitment of β-catenin to the sites of cell-cell junctions (Fig. 2A) and also constructed cell-cell junctions of normal height, indistinguishable from WT E-cad cells (Fig. 2C-D).

Using the quantitative assay for *L. monocytogenes* spread described above, we found a 20% reduction of *L. monocytogenes* focus size in the K738R, K816R E-cad background as compared to the WT E-cad background (Fig. 2B). This observation is consistent with the hypothesis that the JMD of E-cadherin contributes to bacterial cell-to-cell spread in a way that is dependent on E-cadherin ubiquitination by hakai. Because ubiquitination at K738 and K816 is known to induce E-cadherin endocytosis (Fujita et al., 2002), we further hypothesized that E-cadherin could contribute to *L. monocytogenes* cell-to-cell spread by promoting protrusion uptake, a role that is consistent with our demonstration that E-cadherin contributes to bacterial spread at the recipient side of cell-cell contacts. In this scenario, *L. monocytogenes* can be transferred from donor to recipient cell concurrently with E-cadherin endocytosis.

### Clathrin-Mediated Endocytosis of E-cadherin Does Not Promote *L. monocytogenes* Cell-to-Cell Spread

The best-characterized mechanism through which E-cadherin is internalized is clathrin-mediated endocytosis (CME), where adaptor proteins such as Numb and AP-2 bind to E-cadherin to bring about its clathrin-dependent internalization (Le et al., 1999). This pathway seemed a promising candidate, as binding of the protein p120-catenin at the cadherin JMD is known to inhibit endocytosis by obstructing endocytic motifs that permit CME (Miyashita and Ozawa, 2007) and preventing cadherin clustering at clathrin-coated pits (Chiasson et al., 2009). However, clathrin heavy chain and AP-2 were not identified as genes affecting *L. monocytogenes* cell-to-cell spread in an RNAi screen for genes involved in spread (Sanderlin et al., 2019) and clathrin was absent from invaginations made by donor cell protrusions containing *L. monocytogenes* (Dhanda et al., 2020). In accordance with these prior results, we did not observe a reduction of focus size when *L. monocytogenes* propagated through A431D cells treated with pitstop 2, an inhibitor of CME (Dutta et al., 2012). Instead, we observed a small, 10% increase in focus size in the WT E-cad cells and no change in the Δcyto E-cad, Δβ E-cad, or K738R, K816R E-cad cells (Fig. 3A). To confirm that pitstop 2 was inhibiting CME in A431D cells, we quantified uptake of transferrin, a standard target of CME (Harding et al., 1983) and found that it was reduced upon addition of pitstop 2 (Suppl. Fig. S3A). Next, we used an antibody-based surface accessibility assay to measure what proportion of total E-cadherin is available for antibody binding at the cell surface in the presence and absence of pitstop 2. We observed a modest but significant increase in surface E-cadherin in WT E-cad A431D cells upon treatment with pitstop 2, as would be expected if the drug causes a defect in E-cadherin internalization (Suppl. Fig. S3B). These results confirm that CME contributes to E-cadherin internalization in A431D cells, and that pitstop 2 effectively disrupts CME under our assay conditions, but suggest that CME is not a major pathway contributing to *L. monocytogenes* cell-to-cell spread.

**Figure 3:**
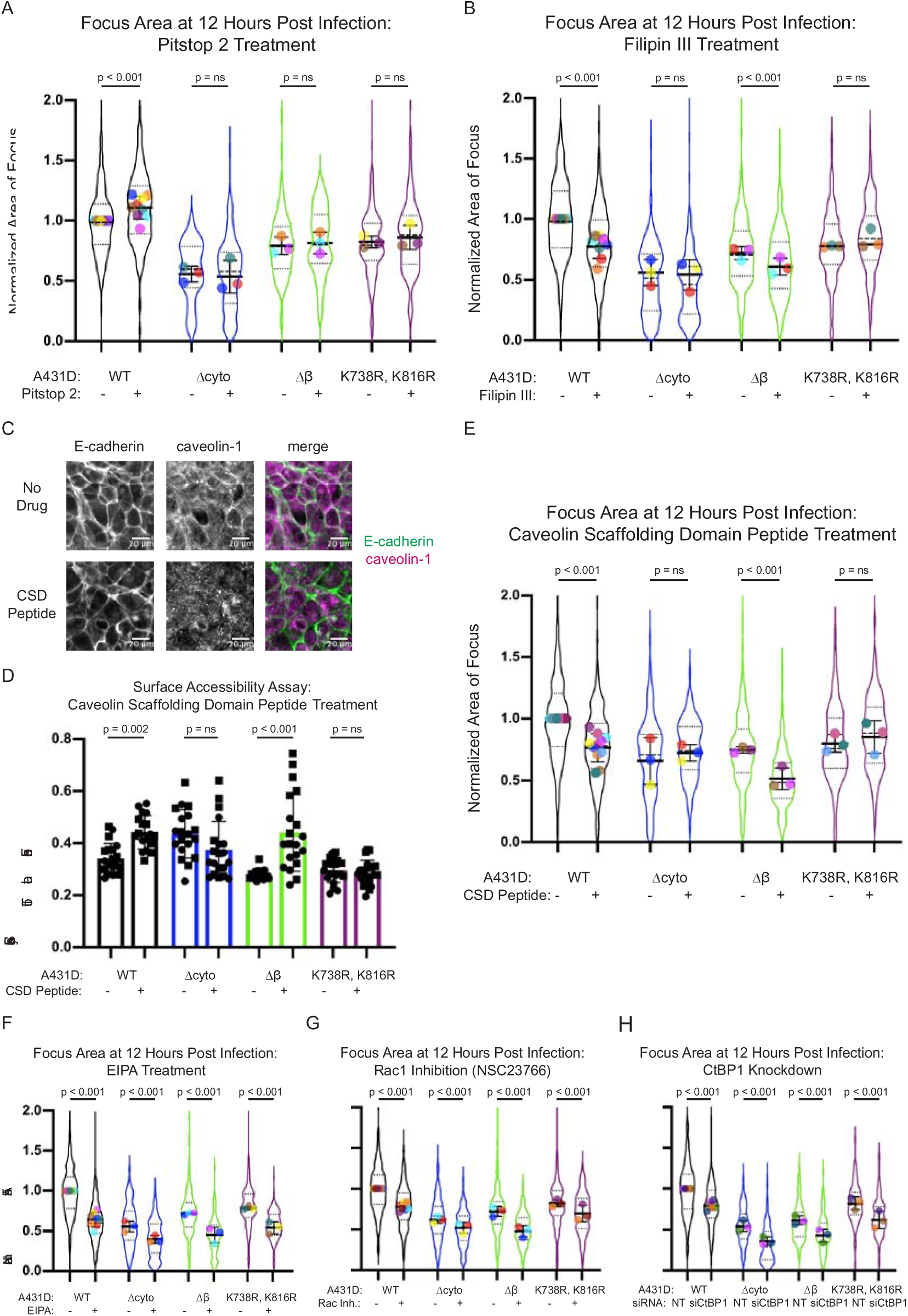
Inhibiting caveolin-mediated E-cadherin internalization and macropinocytosis reduces efficiency of *L. monocytogenes* cell-to-cell spread. **A.** Focus area for A431D cells expressing WT, Δcyto, Δβ and K738R, K816R E-cadherin with and without pitstop 2 treatment. Each colored dot represents a separate experimental day, and all conditions were normalized to the average focus size for the control infection (untreated WT-E-cad host cells) performed on the same day. Between 3 and 9 independent experiments were imaged for each condition, and the number of individual foci imaged for each condition from left to right was 563, 429, 109, 166, 112, 85, 199 and 130. Violin plots show the full range of focus sizes, thick black line shows the mean, whiskers show the standard deviation, dashed line shows the median, and dotted lines the first and third quartiles. P values were determined using the linear mixed-effects model. **B.** Focus area measured as in part A, with and without treatment with filipin III. Between 3 and 8 independent experiments were imaged for each condition and the number of individual foci imaged for each condition from left to right was 433, 375, 185, 177, 259, 217, 159 and 97. P values were determined using the linear mixed-effects model. **C.** Immunofluorescence images showing mislocalization of caveolin-2 upon treatment of WT E-cad A431D cells with CSD peptide. **D.** Measurement of the fraction of E-cadherin that is surface-exposed for A431D cells expressing WT, Δcyto, Δβ and K738R, K816R E-cadherin , with and without treatment with CSD peptide. Circles and squares represent fields of view imaged on two separate coverslips. Mean ± SD is shown on the graph, and the p values were determined using the Wilcoxon rank-sum test. **E.** Focus area measured as in part A, with and without treatment with CSD Peptide. Between 3 and 10 independent experiments were imaged for each condition and the number of individual foci imaged for each condition from left to right was 568, 425, 98, 71, 257, 143, 94 and 73. P values were determined using the linear mixed-effects model. **F.** Focus area measured as in part A, with and without treatment with EIPA. Between 3 and 8 independent experiments were imaged for each condition and the number of individual foci imaged for each condition from left to right was 493, 460. 182, 194, 158, 113, 210 and 103. P values were determined using the linear mixed-effects model. **G.** Focus area measured as in part A, with and without treatment with NSC23766. Between 3 and 7 independent experiments were imaged for each condition and the number of individual foci imaged for each condition from left to right was 458, 491. 235, 217, 211, 159, 199 and 137. P values were determined using the linear mixed-effects model. **H.** Focus area measured as in part A, upon siRNA knockdown of CtBP1 as compared to a non-targeting control (NT). Between 3 and 6 independent experiments were imaged for each condition and the number of individual foci imaged for each condition from left to right was 256, 207. 114, 139, 80, 85, 105 and 110. P values were determined using the linear mixed-effects model.

### *L. monocytogenes* Spreads from Cell to Cell Concurrent with the Caveolin-Mediated Internalization of E-cadherin

An alternative pathway that has been shown to contribute to E-cadherin internalization and turnover in epithelial cells is caveolin-mediated uptake (Lu et al., 2003, Orlichenko et al., 2009, Akhtar and Hotchin, 2001). To assess whether *L. monocytogenes* might use caveolin-mediated E-cadherin internalization to spread from cell to cell, we quantified spread efficiency in A431D cells treated with filipin III, a cholesterol sequestering reagent (Behnke et al., 1984) (Fig. 3B). Focus size was reduced in comparison to untreated cells in the WT E-cad and Δβ E-cad background but not in A431D cells expressing Δcyto E-cad or K738R, K816R E-cad.

As filipin III alters membrane fluidity (Zhang et al., 2019), it can also disrupt endocytic pathways that do not involve caveolin (Hao et al., 2004) and so we sought to confirm our results using the more specific approach of treating A431D cells with the caveolin-scaffolding domain peptide (CSD peptide). This peptide competes with endogenous caveolin-1 to disrupt its interaction with binding partners (Bucci et al., 2000). As the N-terminal cytoplasmic domain of caveolin-1 immunoprecipitates with E-cadherin (Galbiati et al., 2000), the two proteins are likely to associate *in vivo*. In WT E-cad A431D cells treated with vehicle control, caveolin-1 colocalized with E-cadherin at cell boundaries (Fig. 3C). Upon addition of CSD peptide, E-cadherin remained at the cell junctions but caveolin-1 was mislocalized (Fig. 3C), confirming that the peptide interfered with the association between the two proteins. Using the surface accessibility assay, we also demonstrated that CSD peptide reduced E-cadherin internalization in the WT E-cad and Δβ E-cad backgrounds but not in cells expressing Δcyto E-cadherin or K738R, K816R E-cad (Fig. 3D). In addition to suggesting that an interaction between caveolin-1’s scaffolding domain and E-cadherin’s JMD is required for caveolin-mediated E-cadherin internalization, this result indicates that internalization of E-cadherin after ubiquitination by hakai occurs through a caveolin-mediated process.

Using our quantitative assay to measure the efficiency of *L. monocytogenes* cell-to-cell spread, we found that focus size was diminished in WT E-cad and Δβ E-cad cells that were treated with CSD peptide, but remained unaltered in CSD-treated Δcyto E-cad and K738R, K816R E-cad cells (Fig. 3E). This result suggests that caveolin-mediated E-cadherin internalization contributes to *L. monocytogenes* cell-to-cell spread, in a manner that is dependent on ubiquitination of the E-cadherin JMD by hakai.

### Protrusions Containing *L. monocytogenes* Can Be Taken Up By the Recipient Cell Through Macropinocytosis

A third mechanism through which E-cadherin may be internalized is macropinocytosis (Bryant et al., 2007). When we treated A431D cells with 5-[N-ethyl-N-isopropyl] amiloride (EIPA) and Rac1 Inhibitor (NSC23766), inhibitors of macropinocytosis (Koivusalo et al., 2010, Bryant et al., 2007), we observed a reduction of focus size, regardless of E-cadherin background (Fig. 3F, 3G). As an independent approach to measure the role of macropinocytosis in *L. monocytogenes* intercellular spread, we used siRNA knockdown of CtBP1 (Suppl. Fig. S3D), a protein that is required for macropinosome scission (Liberali et al., 2008). Once again, we observed a decrease in efficiency of *L. monocytogenes* cell-to-cell spread after CtBP1 knockdown in A431D cells expressing any of our four E-cadherin constructs (Fig. 3H). These results led us to conclude that macropinocytosis is an additional mechanism through which *L. monocytogenes* spreads from cell to cell, but this mechanism is not dependent on any specific protein interactions with the cytoplasmic domain of E-cadherin.

To confirm that these drug treatments were blocking macropinocytosis under our experimental conditions, we monitored nonspecific macropinocytic uptake of fluorescent dextran from the cell culture medium, and found it to be substantially reduced upon treatment of A431D cells with EIPA and NSC23766 (Suppl. Fig. S3E) as well as after siRNA knockdown of CtBP1 (Suppl. Fig. S3F). As an additional control, we used the surface accessibility assay to demonstrate that EIPA and NSC23766 increased the proportion of E-cadherin remaining on the cell surface (Suppl. Fig. S3G, S3H), as expected if macropinocytosis normally contributes to E-cadherin internalization and turnover in these cells. However, none of these treatments affected the average number of *L. monocytogenes* present in A431D cells in the first 3 to 5 hours post-infection, as determined using a mutant strain of bacteria incapable of actin-based cell-to-cell spread (Suppl. Fig. S3I-M). These results support the conclusion that macropinocytosis contributes to *L. monocytogenes* cell-to-cell spread, but is not involved in earlier stages of bacterial invasion or replication.

### Epidermal Growth Factor (EGF) Treatment Enhances Caveolin-Mediated Internalization of E-cadherin and Promotes *L. monocytogenes* Cell-to-Cell Spread

So far we have shown that inhibiting ubiquitin-dependent caveolin-mediated internalization of E-cadherin and inhibiting macropinocytosis both independently decrease the efficiency of *L. monocytogenes* cell-to-cell spread. We were therefore curious whether stimulation (rather than inhibition) of these pathways might actually enhance the efficiency of bacterial spread. As the growth factor EGF can stimulate both macropinocytosis and caveolin-mediated internalization of E-cadherin (Lu et al., 2003, Bryant et al., 2007), we sought to measure bacterial spread efficiency upon treatment of A431D cells with EGF.

EGF stimulation of macropinocytosis in A431 cells is usually observed in the absence of serum (Hamasaki et al., 2004) but as serum is present in our standard infection assay, it was first necessary for us to measure the effect of EGF on macropinocytosis when we replicated the conditions of our infection assay. Under these conditions, we found no increase in internalized dextran when EGF was added to A431D cells regardless of E-cadherin background (Fig. 4A), suggesting that EGF does not further stimulate macropinocytosis in the presence of serum in A431D cells. Measuring the relative amount of E-cadherin present on the cell surface before and after EGF stimulation using the surface accessibility assay described above, we found that EGF increased E-cadherin internalization in WT E-cad and Δβ E-cad cells but did not alter levels of surface E-cadherin in Δcyto E-cad and K738R, K816R E-cad cells (Suppl. Fig. S4A). Taken together, these results are most consistent with the proposition that EGF stimulates caveolin-mediated internalization of E-cadherin in A431D cells, but does not substantially affect macropinocytosis, at least not under these experimental conditions.

**Figure 4:**
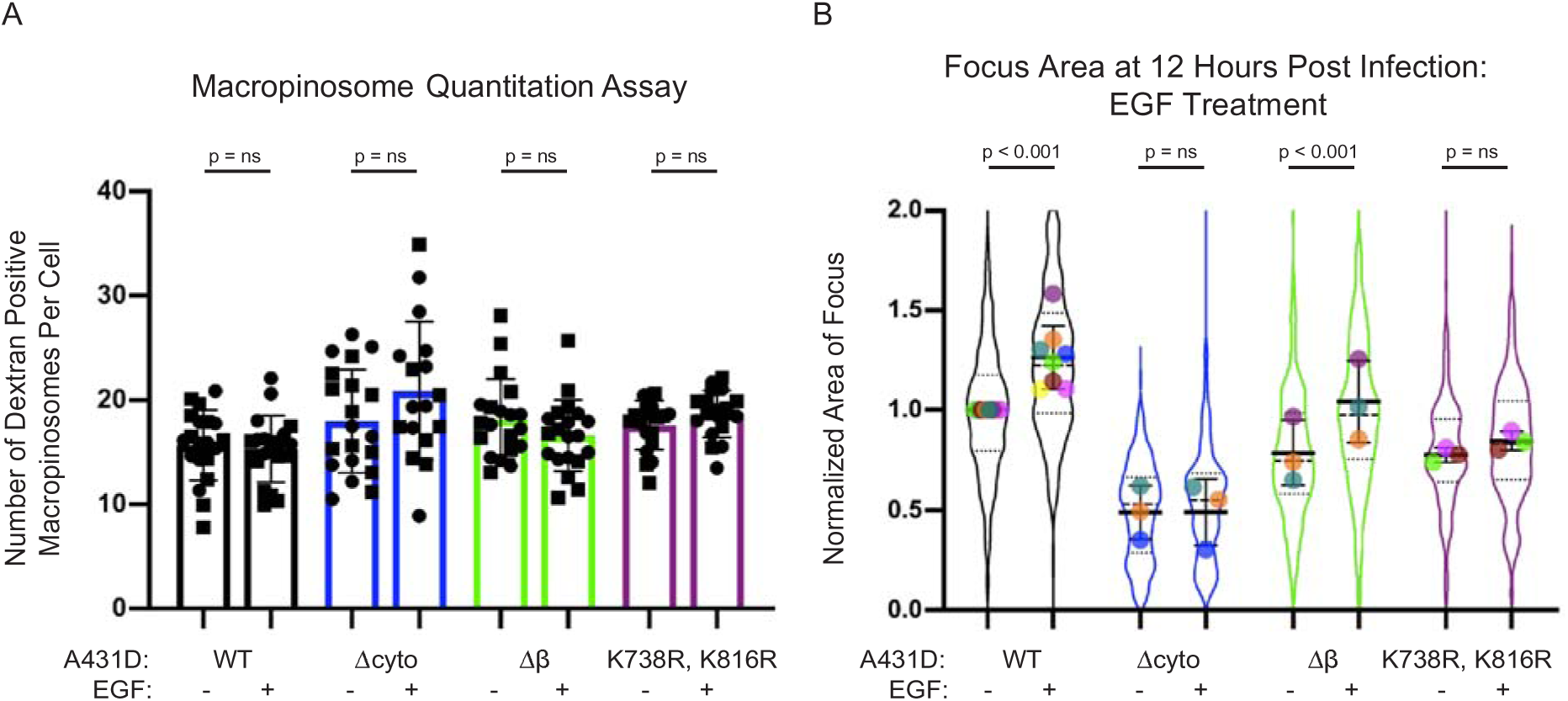
Stimulating caveolin-mediated E-cadherin internalization with EGF enhances *L. monocytogenes* cell-to-cell spread efficiency. **A.** Measurement of the number of dextran-positive macropinosomes in A431D cells expressing WT, Δcyto, Δβ and K738R, K816R E-cadherin, with and without treatment with EGF. Circles and squares represent fields of view imaged on two separate dishes. Mean ± SD are shown on the graph, and the p values were determined using the Wilcoxon rank-sum test. **B.** Focus area for *L. monocytogenes* infections in A431D cells expressing WT, Δcyto, Δβ and K738R, K816R E-cadherin, with and without treatment with exogenous EGF. Each colored dot represents a separate experimental day, and all conditions were normalized to the average focus size for the control infection (untreated WT-E-cad host cells) performed on the same day. Between 3 and 8 independent experiments were imaged for each condition and the number of individual foci imaged for each condition from left to right is 437, 496, 183, 188, 194, 204, 224 and 190. Violin plots show the full range of focus sizes, thick black line shows the mean, whiskers show the standard deviation, dashed line shows the median, and dotted lines the first and third quartiles. P values were determined using the linear mixed-effects model.

When we treated A431D cells with EGF in the presence of serum, and then infected them with *L. monocytogenes*, we observed a modest but statistically significant increase in focus size in WT E-cad and Δβ E-cad A431D cells but not in Δcyto E-cad A431D or K738R, K816R E-cad cells (Fig. 4B). This EGF-dependent increase in focus size is abrogated by treatment with the EGFR inhibitor gefitinib (Lynch et al., 2004), but gefitinib has no effect on focus size in the absence of exogenous EGF (Suppl. Fig. S4B). Finally, we confirmed that EGF treatment does not alter the number of bacteria in foci at early stages of infection (Suppl. Fig. S4C). Given the evidence described above that a major effect of EGF stimulation in these cells is to promote caveolin-mediated uptake of E-cadherin, these results further confirm our central conclusion that this particular pathway for E-cadherin trafficking contributes to the efficiency of *L. monocytogenes* cell-to-cell spread.

## DISCUSSION

Our results support the hypothesis that caveolin-mediated uptake of E-cadherin, but not its clathrin-mediated uptake, promotes the engulfment of *L. monocytogenes*-containing protrusions on the recipient side of host cell-cell contacts, thereby enhancing the efficiency of bacterial cell-to-cell spread in epithelial monolayers. Significantly, we made use of a subtle mutational alteration in the juxtamembrane domain of E-cadherin, converting two lysine residues to arginine residues in order to block ubiquitination of the E-cadherin cytoplasmic domain by the ubiquitin ligase hakai (Hartsock & Nelson, 2012), without disrupting the ability of E-cadherin to mediate formation of normal cell-cell junctions or to interact with the actin cytoskeleton via binding to β-catenin. Using this mutant, we found that disruption of cell-to-cell spread by inhibiting caveolin-mediated trafficking can affect the efficiency of bacterial spread only when E-cadherin itself can be ubiquitinated. Prior work has provided evidence for the involvement of E-cadherin in *L. monocytogenes* cell-to-cell spread (Sanderlin et al., 2019) and suggested a role for caveolins and other caveolar components in this process (Sanderlin et al., 2019; Dhandha et al 2020). We confirm and extend this work by suggesting that it is most likely a direct interaction between caveolin and E-cadherin that facilitates this function.

In addition, our results suggest that macropinocytosis is an additional pathway that contributes to the efficiency of *L. monocytogenes* cell-to-cell spread, although this mechanism is not dependent on any particular domain of E-cadherin. While macropinocytosis is known to be a mechanism through which many pathogens enter host cells (Bloomfield and Kay, 2016) and has also been implicated in the intercellular spread of prions (Zeineddine and Yerbury, 2015) and viruses such as HIV-1 (Wang et al., 2008) and coronaviruses (Freeman et al., 2014), to our knowledge, *L. monocytogenes* is the only bacterial pathogen for which macropinocytosis has been shown to contribute to intercellular spread (Fukumatsu et al., 2012).

Finally, it is illuminating to compare *L. monocytogenes* cell-to-cell spread with the mechanism used by *Shigella flexneri*, another intracellular bacterial pathogen that employs actin-based motility to disseminate through an epithelial monolayer (Bernardini et al., 1989). Spread of both pathogens occurs at cell-cell junctions (Robbins et al., 1999, Rajabian et al., 2009, Duncan-Lowey et al., 2020) and loss of E-cadherin reduces efficiency of spread (Fukumatsu et al., 2012, Sanderlin et al., 2019). However, *S. flexneri* targets tricellular junctions, while *L. monocytogenes* does not exhibit a preference for tricellular over bicellular junctions. Moreover, knockdown of tricellulin, which is enriched at tricellular junctions, greatly diminishes *S. flexneri* cell-to-cell spread but has no effect on *L. monocytogenes* spread (Fukumatsu et al., 2012).

Consistent with our results, Fukumatsu et al. also showed that inhibiting macropinocytosis with EIPA reduced efficiency of *L. monocytogenes* cell-to-cell spread, while suppressing clathrin-mediated uptake with phenylarsine oxide did not significantly impair spread in Caco-2 epithelial cells. Additionally, they showed that inhibiting caveolar uptake with the cholesterol depleting agent, methyl-β-cyclodextrin, limited *L. monocytogenes* cell-to-cell spread, consistent with our results using filipin III to deplete cholesterol and using the CSD peptide to interfere with caveolin protein-protein interactions more directly. Significantly, Fukumatsu et al. showed that the impact of these drugs on *S. flexneri* cell-to-cell spread was distinct from that for *L. monocytogenes*; specifically, macropinocytosis did not appear to contribute to spread at all, while inhibiting caveolin-mediated endocytosis had a modest effect and clathrin-mediated endocytosis was found to be the major pathway *S. flexneri* exploits to spread from cell-to-cell (Fukumatsu et al., 2012). These results suggest that the bacteria are not merely carried into neighboring cells during trans-endocytosis events that are occurring constitutively at some basal level even in uninfected cells (Robbins et al., 1999; Generous et al., 2019, Sakurai et al., 2014). Instead, there appears to be some specificity to the choice of trafficking pathway utilized by each pathogen.

## ACKNOWLEDGEMENTS

We thank Cara Gottardi for the generous gift of the human full-length E-cadherin cDNA as well as the parental A431D cell line. We additionally thank Fabian Ortega for generating the WT, Δcyto and Δβ E-cad A431D cell lines. Furthermore, we thank Fabian Ortega, Matthew Footer, Nathan Belliveau and Gemini Skariah for experimental support as well as members of the Theriot lab for thought-provoking discussion. Effie Bastounis, Matthew Footer, and Fabian Ortega provided valuable feedback on the manuscript. This work was supported by NIAID R37 AI063929-23 (J.A.T.), HHMI (J.A.T.) and the NIH training grant T32 GM 8294-28 (P.R.).

This article is subject to HHMI’s Open Access to Publications policy. HHMI lab heads have previously granted a nonexclusive CC BY 4.0 license to the public and a sublicensable license to HHMI in their research articles. Pursuant to those licenses, the author-accepted manuscript of this article can be made freely available under a CC BY 4.0 license immediately upon publication.

## AUTHOR CONTRIBUTIONS

Conceptualization, P.R. and J.A.T.; Methodology, P.R., M.S., and J.A.T.; Software, P.R.; Validation, P.R.; Formal Analysis, P.R.; Writing – Original Draft, P.R.; Writing – Review & Editing, P.R., M.S., and J.A.T.; Supervision, J.A.T.

## MATERIALS AND METHODS

### Bacterial Strains and Growth Conditions

The WT 10403S strain of *Listeria monocytogenes* was used in this study. Cell-to-cell spread experiments were conducted using WT *L. monocytogenes* that expressed mTagRFP under control of the ActA promoter, so that bacteria would only fluoresce once expression of ActA was induced in the host cell cytosol. This strain was created as previously described (Lauer et al., 2002, Ortega et al., 2017). Briefly, the plasmid pMP74RFP was transformed into *Escherichia coli* SM10 λpir and transferred to *L. monocytogenes* by conjugation, after which the plasmid was integrated into the tRNA^ARG^ locus of the bacterial chromosome. The ΔActA *L. monocytogenes* strain was created by performing an in-frame deletion in the *actA* open reading frame of mTagRFP-expressing WT *L. monocytogenes*.

Before infection experiments, bacteria were streaked from a glycerol stock onto BHI plates supplemented with 200 μg/mL streptomycin and 7.5 μg/mL chloramphenicol. After 48 hours of incubation at 37°C, bacteria were added to 2mL BHI cultures containing the same antibiotics. These cultures were grown overnight in the dark at room temperature without agitation. When the bacteria reached an OD_600_ of 0.8, they were washed twice with PBS and added to A431D cells as described in the infection assay section below.

### Mammalian Cell Culture

A431D human epidermoid carcinoma cells (Lewis et al., 1997) (gift from Cara Gottardi, Northwestern University) were cultured in DMEM supplemented with high glucose (4.5g/L) (Thermofisher; 11965092), 10% fetal bovine serum (FBS) (GemBio; 900-108) and antibiotic-antimycotic (Thermofisher; 15240096). Geneticin (Thermofisher; 10131035), at a concentration of 800μg/mL, was added to media that was used to grow all retrovirally transduced cell lines (WT, Δcyto, Δβ and K738R, K816R E-cad A431D cells).

### Retroviral Transduction of E-cadherin Constructs

As described previously (Ortega et al., 2017), various E-cadherin constructs were derived using the full-length human E-cadherin sequence from the pcDNA3 human E-cadherin plasmid (a gift from Cara Gottardi’s lab at Northwestern University). Δcyto E-cadherin and Δβ E-cadherin constructs were created by deleting amino acids 731-882 and 810-882 respectively from the coding sequence for full-length E-cadherin, while the K738R, K816R E-cadherin construct was generated by mutating two lysines at positions 738 and 816 to arginines. The WT, Δcyto and Δβ E-cadherin constructs were cloned into pLNCX2 Retroviral Vector (Clonetech) by Epoch Life Sciences Inc. GP2-293 cells (Clonetech) were transfected with 10μg of one of the E-cadherin constructs as well as an amphotropic packaging vector (Clonetech). The virus was concentrated 24 hours post transfection and filtered through a 0.45 μm cellulose acetate filter 48 hours post transfection. It was then added to A431D cells along with 8 μg/mL polybrene (Thermofisher; TR1003G) in the transduction step. Transduced A431D cells were selected for using media (high glucose DMEM, 10% FBS, antibiotic-antimycotic (abam)) supplemented with 800 μg/mL of the selective antibiotic geneticin.

### siRNA Transfection of A431D Cells

A431D cells at a density of 2×10^5^ cells/mL in serum and antibiotic-free media were transfected with a final concentration of 20 nM of the selected siRNA in a total reaction volume of 2 mL using lipofectamine RNAiMAX (Invitrogen; 13778075). Transfected cells were seeded onto a fibronectin-coated 12-well glass-bottom plate (Cellvis; P12-1.5H-N) and the transfection mix was replaced with full media 8 hours post transfection. ON-Targetplus siRNA Reagents that used a pool of siRNAs to target our genes of interest were purchased from Dharmacon for use in these experiments. CtBP1was the specific gene that was knocked down. To demonstrate that siRNAs were able to enter host cells, cells were transfected with siGLO, which turns cells fluorescent 24 hours post-transfection. CtBP1 was knocked down between three and six separate times in each E-cadherin expressing A431D cell line.

### Realtime-qPCR

Knockdown of CtBP1 was confirmed using Realtime-qPCR (RT-qPCR). A431D cells treated with siCtBP1 or the non-targeting control were detached from the glass-bottom plate with 0.25% trypsin-EDTA (Thermofisher; 25200056) 72 hours post transfection. The cells were then centrifuged at 300g for 5 minutes and lysed using buffer RLT provided in the RNAeasy Plus Micro Kit (Qiagen; 74034). After the lysate was homogenized using the QIAshredder Kit (Qiagen; 79656), mRNA was extracted using the RNeasy Plus Micro Kit and eluted in 30μL of RNAse-free water. Between 2-3 μg of mRNA from one to two replicates for each condition was provided to Arraystar Inc. for quantitation of gene expression levels using RT-qPCR.

cDNA was synthesized from the mRNA provided using SuperScript III Reverse Transcriptase (Invitrogen; 18080085). Then, RT-qPCR was performed using SuperArray PCR Master Mix to quantify expression of the gene of interest, CtBP1, and a housekeeping gene, β-actin. Primers used to amplify these genes during the course of the PCR reaction are as follows:

CtBP1 Forward: 5’ CACCACCTCATCAACGACTTCA 3’
CtBP1 Reverse: 5’ TGCCTGCTCGCTGTACCAT 3’
β-actin Forward: 5’ GTGGCCGAGGACTTTGATTG 3’
Reverse: 5’ CCTGTAACAACGCATCTCATATT 3’

Concentration of the target and housekeeping gene in each PCR product was determined using Rotor-Gene Real-Time Analysis Software 6.0. The relative amount of CtBP1 DNA in each sample was determined by dividing CtBP1 concentration by the concentration of the housekeeping gene. Then, the relative amount of CtBP1 in each condition was normalized to the mean of the relative amount of CtBP1 in the two replicates of WT E-cad A431D cells treated with non-targeting siRNA.

### Infection Assay

To conduct infection assays, A431D cells were seeded on cleaned 18 mm glass coverslips that were coated with 1μg/mL fibronectin at a density of 4×10^5^ cells/mL and were allowed to incubate at 37°C for 36 hours. At this point, one mL of the bacterial culture at OD 0.8 was centrifuged at 2000g for four minutes. The BHI media was then aspirated and the pellet was washed twice with PBS before resuspension in DMEM. Host A431D cells were washed with DMEM once before the diluted bacteria was added to them at a multiplicity of infection (MOI) of 200-300 bacteria per cell. Bacteria and host cells were incubated at 37°C for 10 minutes, after which the host cells were washed three times with DMEM to remove non-adherent bacteria. For the cell-to cell spread experiments, host cells were then incubated in DMEM at 37°C for 15 minutes and subsequently incubated for an additional 12 hours in DMEM supplemented with 10% FBS and 20 μg/mL gentamicin, an antibiotic which kills extracellular bacteria to prevent further invasion (Portnoy et al., 1988). Drug treatments included 30 μM pitstop 2 (Aobious Inc; AOB3600), 5 μM caveolin-scaffolding domain peptide (Sigma Aldrich Inc; 219482), 5 μg/mL filipin III (Thermofisher; 62501NB), 100 ng/mL EGF (Sigma Aldrich Inc,. E9644), 10 μM EIPA (Thermofisher; 337810) and 100 μM NSC23766 (Sigma Aldrich Inc,. 553502). All drugs were added to infected cultures at the same time as gentamicin. The effect of each perturbation on efficiency of spread was assessed on at least three experimental days with two coverslips per condition on each day. For the experiments that assessed the effect a particular drug or cell line might have on the rate of bacterial replication, host cells were incubated in DMEM (neat) at 37°C for 30 minutes before addition of gentamicin and drug. Infection was then allowed to proceed for three, four, and five hours before fixation. Coverslips in all categories of experiments were fixed with 4% PFA for 10 minutes at room temperature, permeabilized with 0.1% Triton X-100 (Sigma Aldrich Inc; T9284) for an additional 10 minutes and stained with the nucleus label DAPI (Thermofisher; D1306), in blacking solution for 45 minutes. Coverslips were then mounted onto slides using VECTASHIELD mounting medium (Vector Labs; H-1000) prior to imaging.

To assess whether E-cadherin promotes spread at the recipient side of cell contacts, monolayers of A431D cells expressing either WT E-cadherin or a co-culture of 1:100 WT E-cad: Null E-cad A431D cells were infected for five hours and subsequently fixed and stained. After imaging, the number of bacteria within the WT E-cad donor cell was counted and divided by the number of bacteria in each of the neighboring recipient cells. This value represented the ratio of bacteria that had been transferred from donor to recipient cell in one round of cell-to-cell spread. Data shown in Figure 1G originates from two independent experiments.

In the case of spread between WT E-cad and Null E-cad cells, the WT E-cad cell is assumed to be the donor, as *L. monocytogenes* adheres to E-cadherin on the cell surface prior to invading epithelial cells (Ortega et al., 2017). Time lapse experiments revealed that the cell at the center of the focus was most likely to be the donor in the case of spread between WT E-cad cells (Data not shown).

### Immunostaining to Visualize Membrane and Protein Localization in A431D Cells

A431D cells were seeded on clean fibronectin-coated 25mm coverslips at a density of 7.5×10^5^ cells/mL and incubated at 37°C for 36 hours. They were subsequently fixed and permeabilized as described above. Then, they were blocked with 1% BSA, 22.5 mg/mL glycine in PBST (PBS+0.1% Tween-20) for 30 minutes to prevent nonspecific binding of antibodies. Subsequently, the cells were incubated with the prescribed concentration of primary antibody (for E-cadherin (Abcam; ab1416), β-catenin (Abcam; ab32572), and caveolin-1 (Abcam; ab2910)) in PBST for 1 hour. They were then washed with PBS three times and incubated with secondary antibody: either Alexa Fluor 488 Goat Anti-Rabbit IgG (H+L) antibody (Thermofisher; A-11008) or Alexa Fluor 647 Donkey anti-Rabbit IgG (H+L) secondary antibody (Thermofisher; A-31573) diluted 1:200 in 2% BSA in PBS for 1 hour in the dark. Lastly, the coverslips were washed 3 times with PBS and mounted onto slides using VECTASHIELD mounting medium prior to imaging. These images were collected using a Plan Apo 60x NA 1.27 water immersion objective with a Yokogawa W1 Spinning Disk Confocal with Borealis Upgrade on a Nikon Eclipse Ti2 inverted microscope with a 50μm disk pattern (Andor). Micromanager (Edelstein et al., 2014) was used to control all microscopy equipment.

Cell membranes of A431D cells were labeled using 5μg/mL rhodamine labeled wheat germ agglutinin (Vector Labs; RL-1022) in HBSS (Hank’s balanced salt solution) for 10 minutes at room temperature. After washing with PBS and permeabilizing with 0.1% Triton X-100 as described above, cells were stained for E-cadherin using the protocol previously outlined and were also labeled with DAPI to illuminate the cell’s nucleus as detailed in the infection assay section. Then, coverslips were mounted onto slides and superresolution images were collected using a Nikon Apo TIRF 100X 1.49 NA objective for STORM/SIM using a VisiTech iSIM coupled with an inverted Nikon Eclipse TI2 and a Hamamatsu ORCA/Fusion sCMOS Camera.

### Image Acquisition and Quantitation of Bacterial Focus Size

We acquired confocal images of mTagRFP expressing *L. monocytogenes* as well as DAPI-stained host cell nuclei with approximately 10 slices per infection focus and a z-spacing of 1μm. These images were taken using a Yokogawa W1 Spinning Disk Confocal with Borealis Upgrade on a Nikon Eclipse Ti2 inverted microscope with a 50μm disk pattern (Andor). Furthermore, a Plan Apo 20X 0.95 NA water immersion objective was used to acquire the images, a piezo z-stage (Ludl 96A600) was used to increase the rate of image acquisition between z-frames, and MicroManager v.1.4.23 was used to control all microscopy equipment. Upto fifty bacterial foci were imaged on each coverslip with two replicate coverslips per condition and each experiment repeated on three separate days.

After performing a maximum intensity projection of the confocal images and cropping out the bacterial focus to eliminate extraneous bacteria in Fiji, Matlab_R2018B was used to threshold and binarize the image of the focus. We then generated an alpha shape around each binarized focus using the Matlab function alphaShape with an alpha radius of 30, suppressed all holes within the alpha shape and calculated its area. This was a deviation from the traditional use of the convex hull area as a measure of spread efficiency as the convex hull tends to highly weight outlying bacteria that occupy extreme positions within the focus (Ortega et al., 2019).

### Time-Lapse Microscopy

WT E-cad and Null E-cad A431D cells seeded onto fibronectin-coated glass bottom plates (Cellvis; P12-1.5H-N) were infected with *L. monocytogenes* as outlined above. At five hours post infection, host cell nuclei were stained with Hoechst (Invitrogen; H3570) in FluoroBrite DMEM (Thermofisher; A1896701) with 20 μg/mL gentamicin for 15 minutes at 37°C. After three washes, the cells were immersed in FluoroBrite DMEM with 10% FBS and 20μg/mL gentamicin and were imaged every five minutes until 12 hours post infection using an inverted Nikon Eclipse TI2 with an EMCCD Camera (Andor Technologies) and a 20X 0.75 NA Plan Apo Air Objective. Micromanager was used to operate all microscopy equipment. An environmental chamber was used to maintain the temperature at 37°C and deliver 5% CO_2_ to the cells.

### Surface Accessibility Assay

A431D cells were seeded on a 35 mm glass-bottom dish with a 20mm well (Cellvis; D35-20-1.5N) that was coated with 1μg/mL fibronectin at a density of 6×10^5^ cells/mL and allowed to incubate at 37°C for 36 hours. At this point, cells were treated with DMSO control or drugs such as filipin III, caveolin-scaffolding domain peptide, EGF, EIPA, NSC23766 and pitstop 2 for 45 minutes at 37 °C. Four dishes were utilized for each condition. Of those, two dishes were transferred onto ice, washed with PBS and labeled with mouse monoclonal antibody to E-cadherin (Abcam; ab1416) diluted 1:1000 in PBS for 30 minutes. Still on ice, these cells were washed with PBS, fixed with 4% methanol-free PFA (Thermofisher; 28906) for 10 minutes and then fixed for 10 additional minutes at room temperature. After another wash, A431D cells were stained with Alexa Fluor 488 Goat Anti-Mouse IgG (H+L) antibody (Thermofisher; A-11008) diluted 1:200 for 30 minutes at room temperature and washed with PBS until no extraneous background fluorescence was detected (usually 5 times). The other two dishes used for a condition were similarly transferred onto ice but fixed first using the same method described above and then permeabilized with 0.1% Triton-X 100 at room temperature for 10 minutes. The protocol for staining cells was identical to the non-permeabilized sample except that all steps were conducted at room temperature. Epifluorescence images of 10 fields of view (FOV) per dish were obtained using an inverted Nikon Eclipse TI2 with an EMCCD Camera (Andor Technologies) and a 20X 0.75 NA Plan Apo Air Objective and Micromanager. The average intensity of each FOV in the non-permeabilized sample was divided by the average intensity of a FOV in the corresponding permeabilized sample to obtain a ratio of surface E-cadherin to total E-cadherin.

### Dextran Uptake Assay

To conduct the dextran uptake assay, A431D cells were seeded on a 35 mm glass-bottom dish with a 20 mm well (Cellvis; D35-20-1.5N) that was coated with 1μg/mL fibronectin at a density of 6×10^5^ cells/mL and allowed to incubate at 37°C for 36 hours. At this point, the confluent monolayer of A431D cells were incubated with L-15 (Life Tech Corp; 21083027) in a water bath at 37 °C for a half hour prior to the dextran pulse. This step was conducted to mimic the infection assay by serum-starving cells for about a half hour before macropinocytosis enhancers or inhibitors were added onto the cells. Furthermore, as L-15 lacks phenol red, this step serves the additional purpose of reducing background fluorescence during the image acquisition. 70 kDa fluorescein dextran (Molecular Probes; D1822) was added to L15-10% FBS at a concentration of 1mg/mL along with 100 ng/mL EGF, 10 μM EIPA, 100 μM NSC23766, or no drug for the control. A431D cells were then pulsed with this solution for 10 minutes in a 37 °C water bath and then quickly transferred onto ice to inhibit any further macropinocytosis. Next, the cells were washed with ice cold PBS, fixed with 4% methanol-free PFA for 30 minutes and washed three more times in PBS. To prevent the quenching of fluorescein dextrans in endosomes due to its low pH, all samples were imaged in a solution of 1mM ammonium chloride in PBS. Images were acquired using a Yokogawa W1 Spinning Disk Confocal with Borealis Upgrade on a Nikon Eclipse TI2 inverted microscope with a 50μm disk pattern using a 40X 1.25 NA Plan Apo objective and Micromanager. After background subtraction, thresholding, and binarization, particles that were 0.5-5.0 μm in diameter were filtered out and counted using Fiji. By dividing this number of particles by the number of cells per FOV, we were able to obtain an average number of internalized dextrans per cell in each condition.

### Transferrin Uptake Assay

To conduct the transferrin internalization assay, A431D cells were seeded on a 35 mm glass-bottom dish with a 20 mm well (Cellvis; D35-20-1.5N) that was coated with 1μg/mL fibronectin at a density of 6×10^5^ cells/mL and allowed to incubate at 37°C for 36 hours. Confluent monolayers of A431D cells in 35mm glass dishes were then incubated with L-15 in a water bath at 37 °C for a half hour prior to the transferrin pulse. Transferrin, Alexa Fluor 647 Conjugate (Thermofisher; T23366), was added to L-15 - 10% FBS at a concentration of 50 μg/mL along with 30 μM pitstop 2, 5 μM caveolin-scaffolding domain peptide, 5 μg/mL filipin III, 100 ng/mL EGF, 10 μM EIPA, 100 μM NSC23766, or no drug for the control condition. A431D cells were then pulsed with this solution for 10 minutes in a 37 °C water bath and then quickly transferred onto ice to inhibit any further endocytosis. After washing with ice-cold PBS, the cells were incubated in ascorbate buffer (160mM sodium ascorbate, 40mM ascorbic acid, 1mM MgCl_2_, 1mM CaCl_2_) twice for five minutes each to strip transferrin from the surface of the cell. They were then incubated in ice-cold PBS for five minutes to return the sample to a neutral pH. Next, cells were fixed on ice with 4% methanol-free PFA for 30 minutes before transferrin receptor was labeled with CD71 Transferrin Receptor Antibody (Thermofisher; 14-0719-82) as a primary antibody and Alexa Fluor 488 Goat Anti-Rabbit IgG (H+L) antibody as the secondary. A431D cells were incubated with both antibodies for 30 minutes each and three PBS washes were conducted post-fixation, after addition of the primary and secondary antibodies. Two independent experiments are reported in Figure S4A. Images were acquired and the ratio of internalized labeled transferrin to surface transferrin receptor was obtained by dividing the average fluorescence intensity of the former from the latter.

### Quantification and Statistical Analysis

Statistical parameters and significance as well as the number of data points evaluated for each experiment are reported in the Figures and the Figure Legends.

Normalized focus area in Figures 1E, 2B, 3A, 3B, 3E, 3F, 3G, 3H, 4B, S4B and normalized nuclei count per field of view in Figure S2B were represented as violin plots. In the violin plots, the dashed line denotes the median of the distribution and the dotted lines represent the 25 and 75% quartile of the pooled data from all independent experiments. The mean for each individual experiment is indicated within the violins as a filled circle. P-values for the comparison of cell-to-cell spread efficiency were calculated using the linear mixed-effects model (Gałecki and Burzykowski, 2013), which takes into account the variability introduced by random effects such as the day of experimentation and coverslip on which cells were seeded. As at least three independent experiments were conducted to evaluate whether a perturbation affected the focus area, the threshold for significance was set at 0.01 to account for multiple hypothesis testing.

The results of the surface accessibility assays (Figures 3D, S3B, S3E, S3F, S4A), transferrin internalization assay (Figure S3A) and macropinosome quantitation assay (Figures S3D, 4A) were represented as bar graphs with the midlines signifying the mean of the data points and the whiskers denoting the standard deviation. Ratio of junction length to cell height (Figure 2C) and bacterial count was represented in the same way in Suppl. Figs. S1D, S2A, S3G, S3H, S3I, S3J, S3K, and S4C. The ratio of bacteria in recipient cells to the donor cell in Figure 1G was represented as a beeswarm plot with midlines signifying the mean of the data points and the whiskers denoting the standard deviation. P-values were calculated using the Wilcoxon rank-sum test in GraphPad PRISM8 with a threshold of significance set at 0.05.

## RESOURCE AVAILABILITY

### Lead contact

Requests for reagents and detailed protocols may be directed to Julie A. Theriot.

### Materials availability

Materials developed in this study are available on request to the corresponding author.

### Data and code availability

Data collected and computer codes are available on request to the corresponding author.

## KEY RESOURCES TABLE

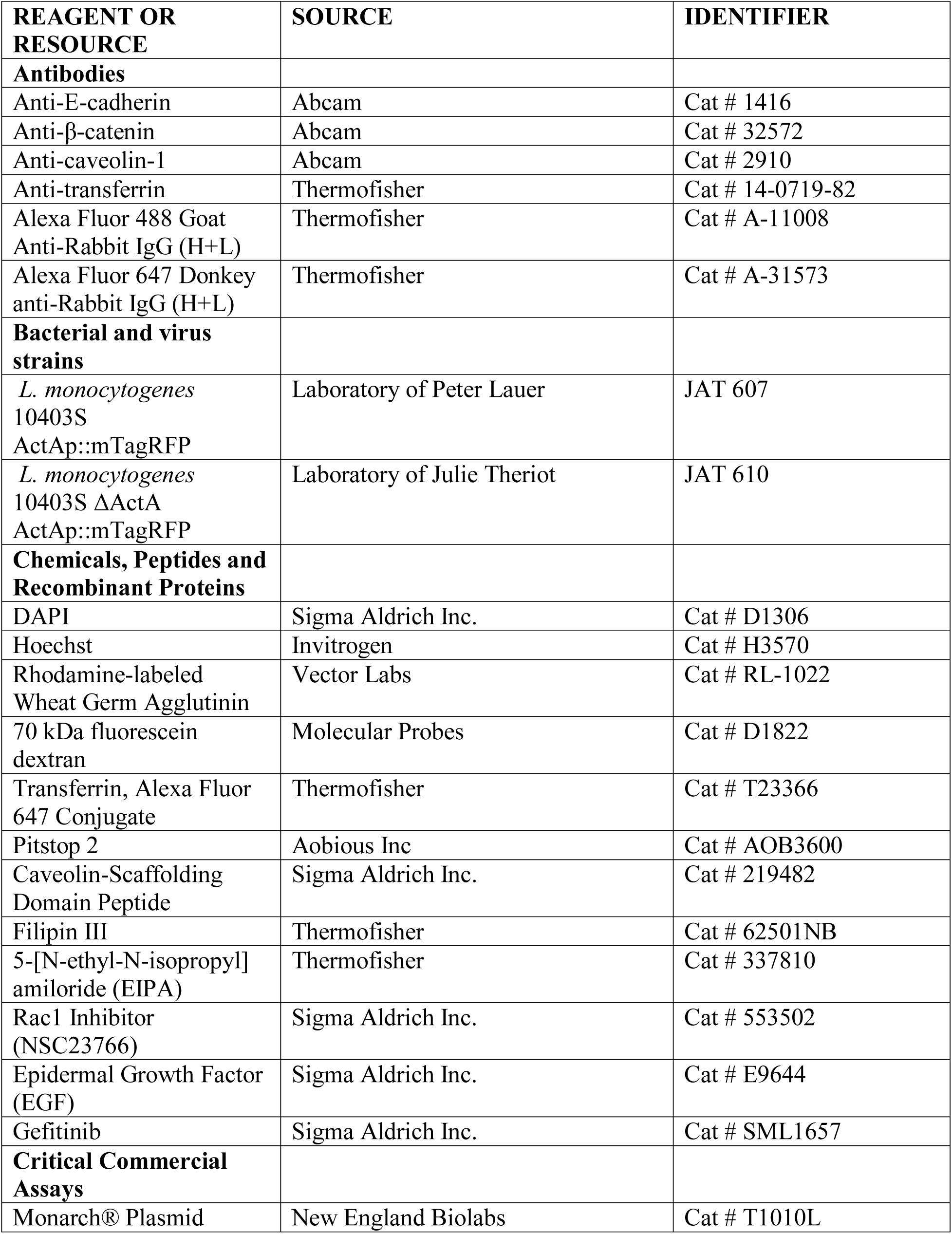
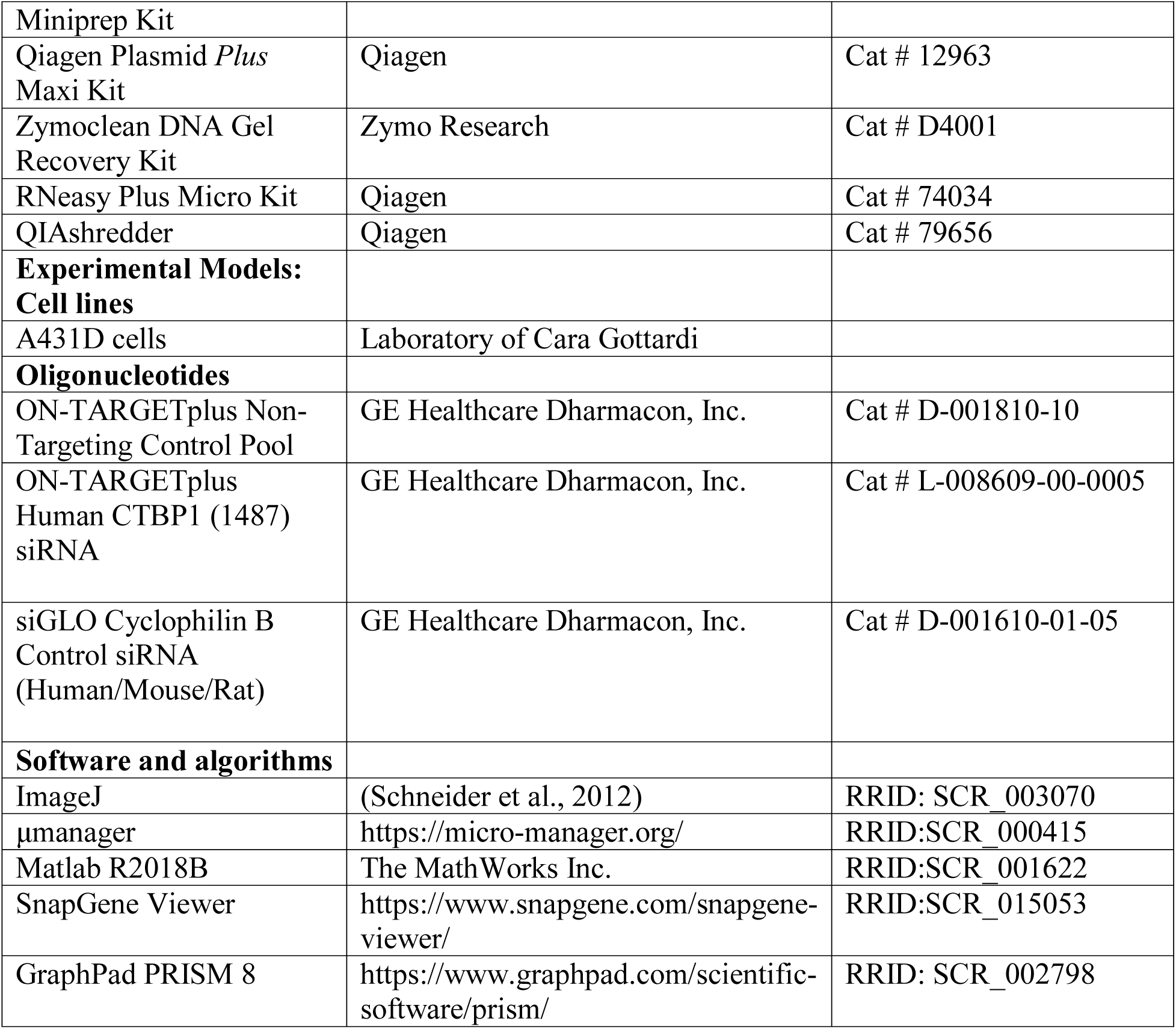

