## Supplemental Figures for "*Listeria monocytogenes* Co-opts Caveolin-Mediated E-cadherin Trafficking and Macropinocytosis for Epithelial Cell-to-Cell Spread"

### SUPPLEMENTAL MATERIALS

Supplemental Figures S1-S4 with legends
Supplemental Movie 1 Legend

#### SUPPLEMENTAL FIGURE S1

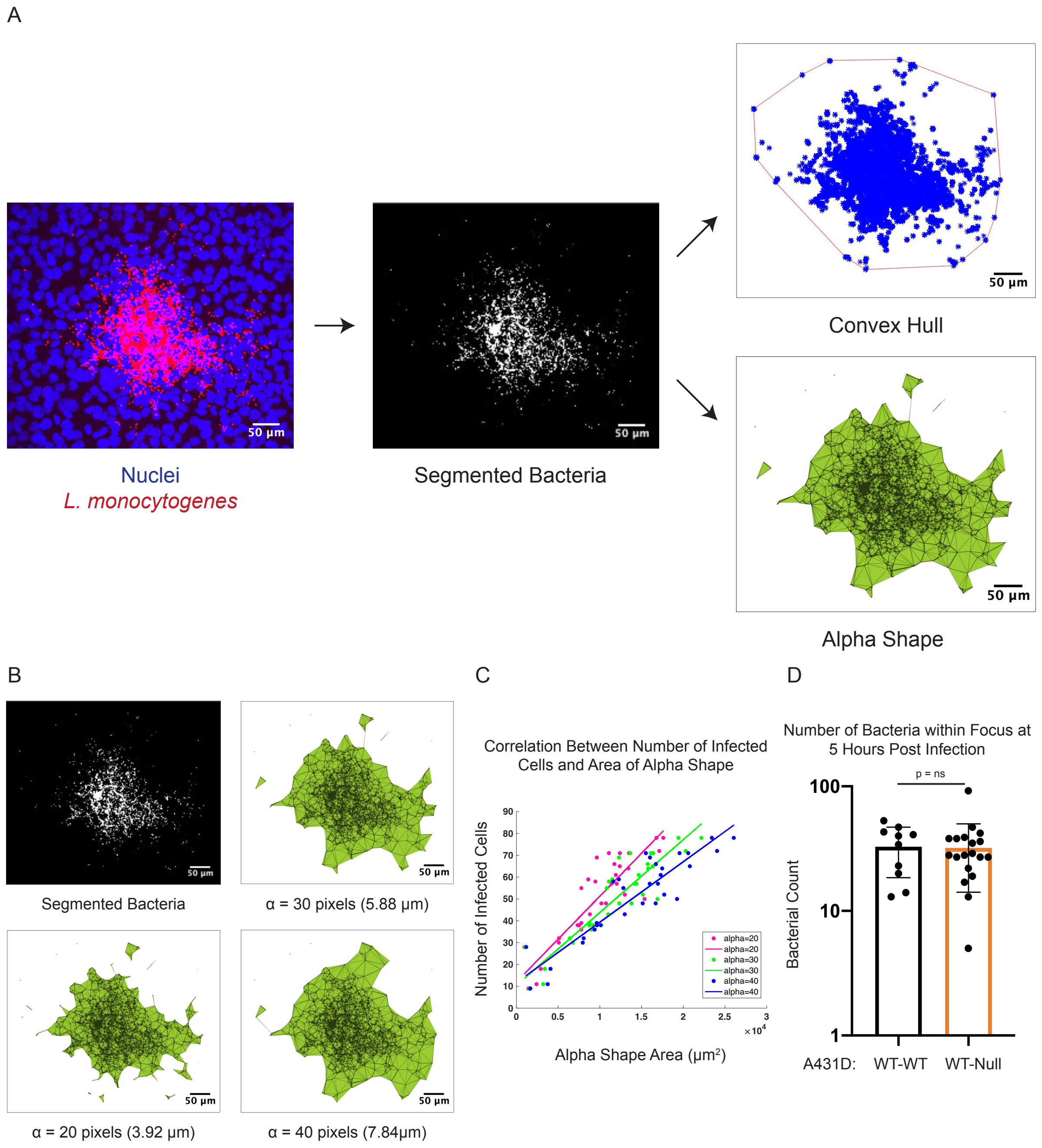

**Supplemental Figure S1: Alpha shapes provide an accurate measurement of *L. monocytogenes* focus size.**

**A.** Maximum intensity projections of *L. monocytogenes* foci are binarized and segmented. Host cell nuclei labeled with DAPI are shown in blue and bacteria expressing mTagRFP are shown in red (left), while binarized bacteria are shown in white (center). Rather than quantifying the area of the convex hull around the focus (top), an alpha shape is generated to circumscribe the focus (bottom) and its area used to represent efficiency of cell-to-cell spread. **B.** Alpha shape representations of one bacterial focus with $\alpha = 20$, 30 or 40 pixels. $\alpha$ determines the tightness with which the alpha shape wraps around the focus, where the alpha shape with $\alpha = 20$ bounds the focus closely and the alpha shape with $\alpha = 40$ forms a looser boundary. **C.** Correlation between alpha shape area and number of infected cells for 30 bacterial foci. Magenta, green and blue dots represent alpha shapes drawn using $\alpha =$ 20, 30, or 40 pixels respectively. **D.** Number of bacteria per focus in WT E-cad and 1:100 WT E-cad to Null E-cad A431D cells. 10 and 19 foci were included for the two conditions. Mean ± SD is shown on the graph, and the p value was determined using the Wilcoxon rank-sum test.

#### SUPPLEMENTAL FIGURE S2

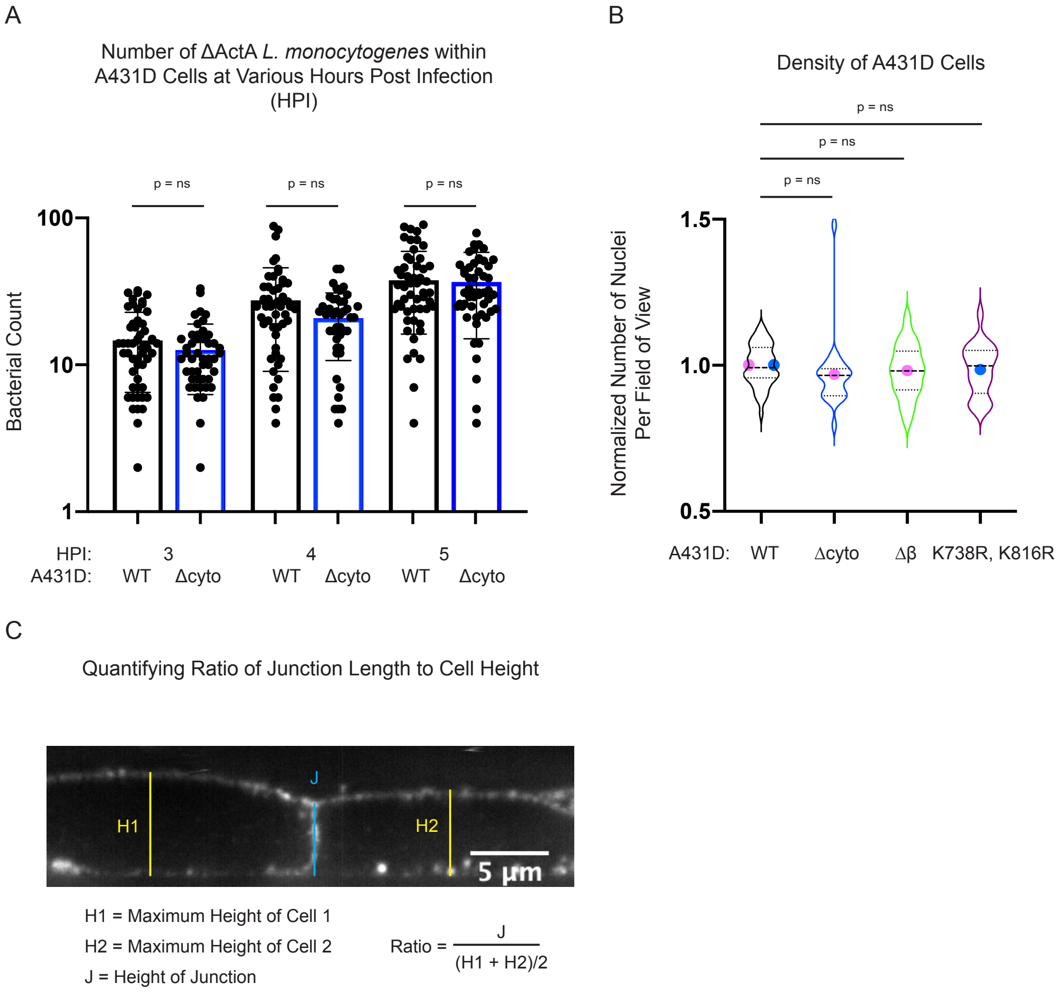


**Supplemental Figure S2: Reduced spread in A431D cells expressing E-cadherin mutants cannot be explained by changes in bacterial proliferation, monolayer confluency, or physical proximity between neighboring cells.**

**A.** Bacterial number for *ΔactA L. monocytogenes* foci in WT E-cad and Δcyto E-cadherin cells at 3, 4, and 5 hours post infection. Between 46 and 52 bacterial foci were analyzed for each condition, mean ± SD is shown on the graph, and the p values were determined using the Wilcoxon rank-sum test. **B.** Number of nuclei per field of view in monolayers of WT, Δcyto, Δβ and K738R, K816R E-cadherin cells normalized to the mean for a field of view of WT E-cadherin cells. Violin plots show the range of normalized nucleus counts within 21 to 42 fields of view per condition with the colored dots representing the mean for two independent days of experimentation. The thick black line shows the mean for the combined data, whiskers show the standard deviation, dashed line shows the median, and dotted lines the first and third quartiles. P-values were calculated using the linear mixed-effects model. **C.** The ratio of junction length to cell height was determined by dividing the height of the junction by the average of the maximal heights of two cells that form the junction. Yellow lines depict the distance from the base of each cell to the highest point on the apical surface, while the blue line depicts the length of the boundary between two cells.

### SUPPLEMENTAL FIGURE S3

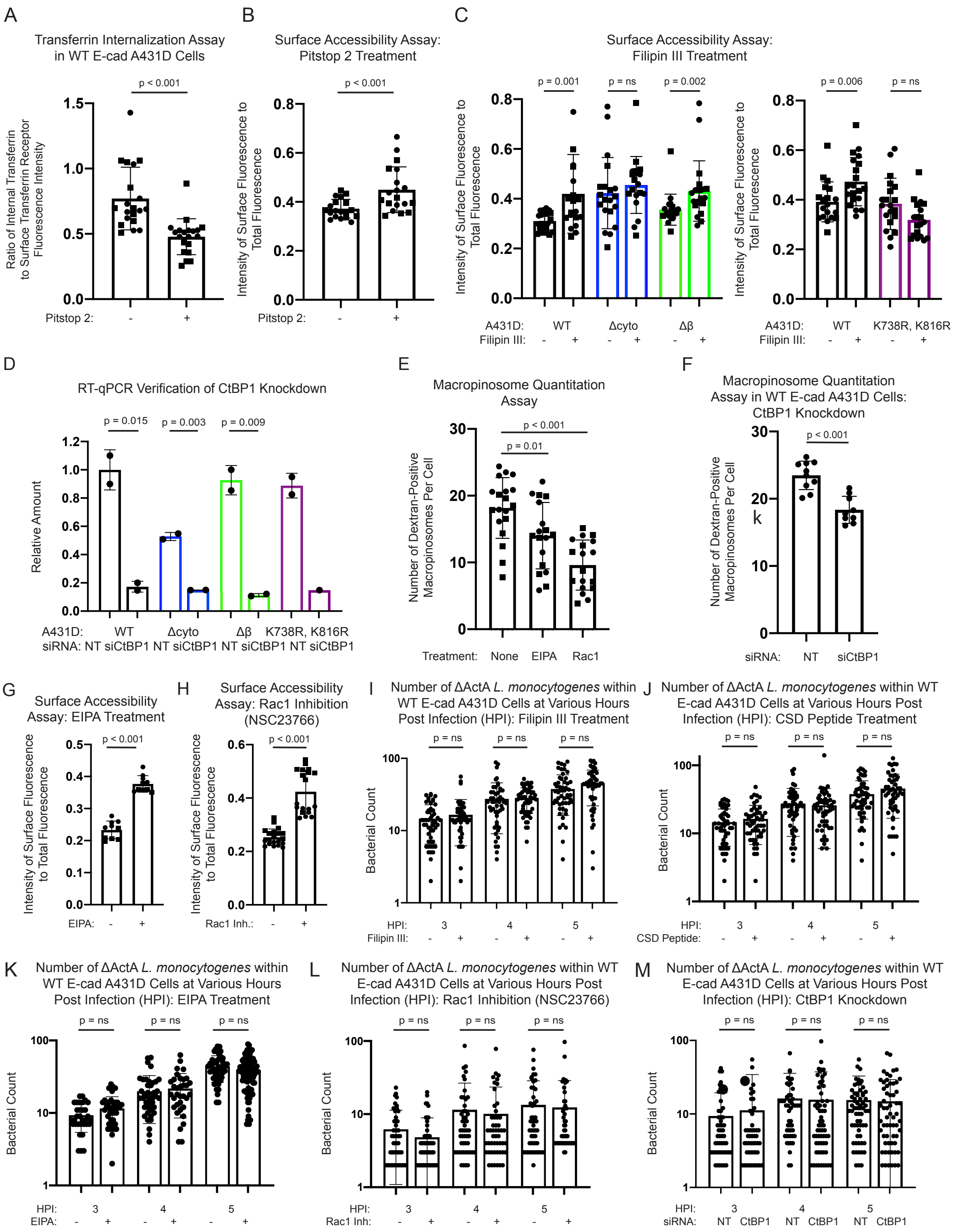

**Supplemental Figure S3: Drug perturbations and CtBP1 knockdown inhibit endocytosis and do not alter bacterial proliferation.**

**A.** Measurement of transferrin uptake efficiency with and without pitstop 2 in WT E-cadherin A431D cells. Circles and squares represent two independent experiments. **B.** Fraction of surface-exposed to total E-cadherin upon treatment of WT E-cad A431D cells with pitstop 2. **C.** Ratio of surface to total E-cadherin upon treatment of WT, Δcyto, Δβ and K738R, K816R E-cad with filipin III as compared to DMSO control. **D.** Relative amount of CtBP1 mRNA in A431D cells treated with siCtBP1 as compared to non-targeting siRNA. Each data point represents expression levels of CtBP1 in cells seeded onto a separate well. Mean ± SD is shown on the graph and p values were determined using the Wilcoxon rank-sum test**. E.** Number of dextran-positive macropinosomes per cell in WT E-cadherin A431D cells treated with EIPA or NSC23766 or in the presence of DMSO control. **F.** Quantification of dextran-positive macropinosomes per cell in WT E-cad A431D cells transfected with non-targeting siRNA or siCtBP1. **G.** Measurement of the fraction of E-cadherin that is surface-exposed in WT E-cad A431D cells with and without EIPA. **H.** Ratio of surface to total E-cadherin upon treatment of WT E-cad A431D cells with NSC23766. **B-C, G-H.** Circles and squares represent fields of view imaged on separate coverslips. Mean ± SD is shown on the graph and p values were determined using the Wilcoxon rank-sum test. **I-M.** Bacterial number for *ΔactA L. monocytogenes* foci in WT E-cad A431D cells treated with filipin III **(I),** CSD Peptide **(J),** EIPA **(K),** NSC23766 **(L)** and siCtBP1**(M)** as compared to control cells at 3, 4, and 5 hours post infection. Mean ± SD is shown on the graph and p values were determined using the Wilcoxon rank-sum test.

### SUPPLEMENTAL FIGURE S4

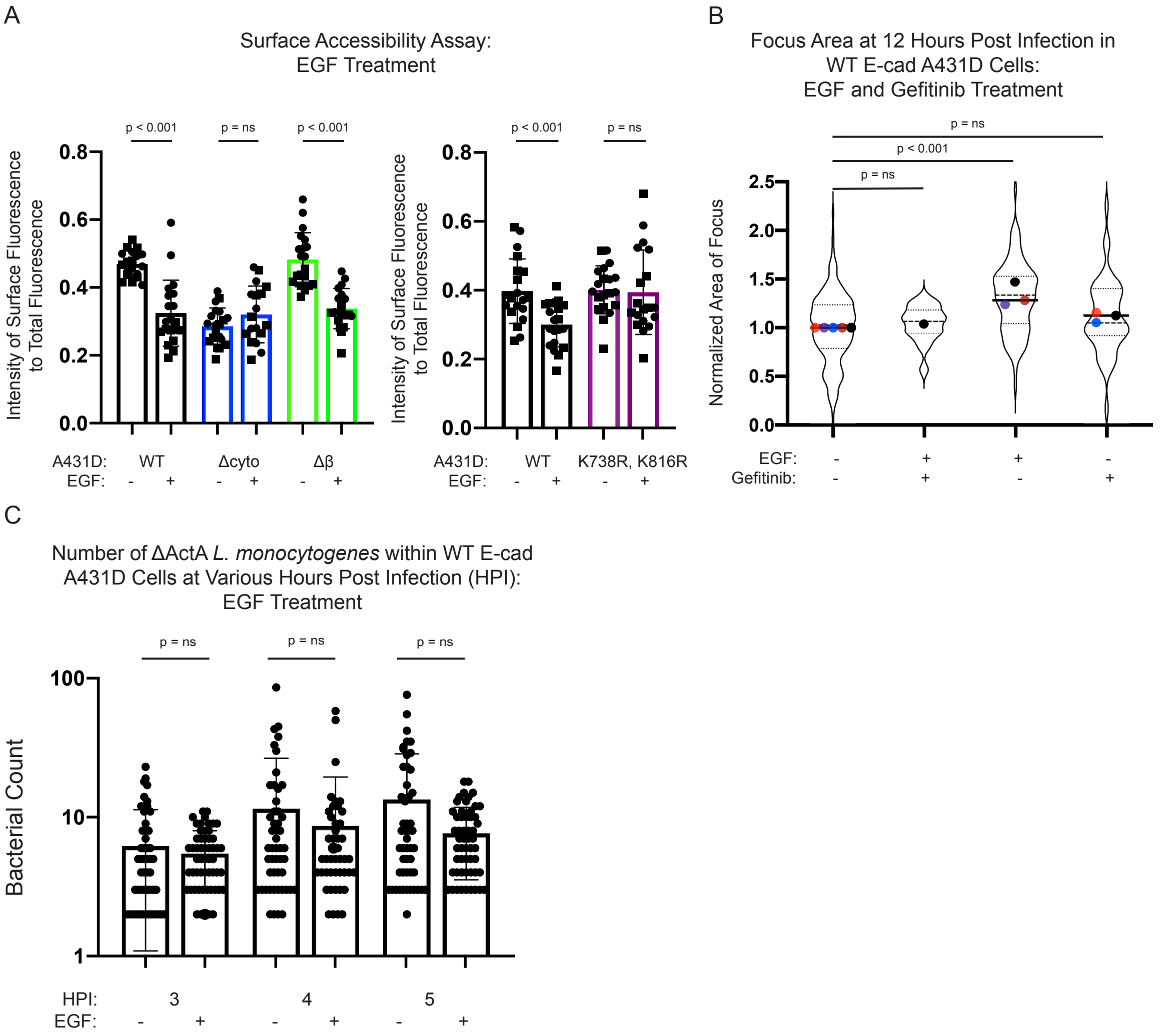


**Supplemental Figure S4: EGF treatment stimulates caveolin-dependent E-cadherin internalization and does not alter bacterial proliferation.**

**A.** Measurement of ratio of surface-exposed E-cadherin to total E-cadherin upon treatment of A431D cells with EGF. Squares and circles represent fields of view imaged on separate coverslips, mean ± SD is shown on the graph, and the p value was determined using the

Wilcoxon rank-sum test. **B.** Focus size for *L. monocytogenes* infection in WT E-cadherin A431D cells which have been treated with gefitinib, gefitinib and EGF or EGF only, normalized to focus size for bacteria spreading through untreated A431D cells. Violin plots show the full range of focus sizes, thick black line shows the mean, whiskers show the standard deviation, dashed line shows the median, and dotted lines the first and third quartiles. Between 1 and 5 independent experiments were imaged for each condition and the number of individual foci imaged for each condition from left to right is 166, 17, 95 and 46. P values were determined using the linear mixed effects model. **C.** Bacterial number per *ΔactA L. monocytogenes* foci in WT E-cad A431D cells treated with EGF contain similar bacterial loads at 3, 4, and 5 hours post infection as untreated cells. Between 46 and 52 bacterial foci were analyzed for each condition. Mean ± SD is shown on the graph and the p values were determined using the Wilcoxon rank-sum test.

**Supplemental Movie 1: Efficiency of *L. monocytogenes* cell-to-cell spread is reduced in 1:100 WT:Null E-cad A431D cells as compared to WT A431D cells**

Time-lapse recording of *L. monocytogenes* spreading through monolayers of epithelial cells that either contain cells that all express full-length E-cadherin (left) or contain a 1:100 mixture of full-length E-cadherin-expressing cells with A431D cells that lack E-cadherin. Bacteria express mTagRFP and are shown in red, while host cell nuclei are labeled with Hoechst and are shown in blue. Images were collected at 3 minute time intervals beginning at 6 hours post infection and continuing for 4 hours. Time shown in hours post infection.
